# Pulsed low-dose-rate radiation (PLDR) reduces the tumor-promoting responses induced by conventional chemoradiation in pancreatic cancer-associated fibroblasts

**DOI:** 10.1101/2024.01.13.575510

**Authors:** Janusz Franco-Barraza, Tiffany Luong, Jessica K. Wong, Kristopher Raghavan, Elizabeth Handorf, Débora B. Vendramini-Costa, Ralph Francescone, Jaye C. Gardiner, Joshua E. Meyer, Edna Cukierman

**Author notes:** Members of the Marvin & Concetta Greenberg Pancreatic Cancer Institute.

## Abstract

Pancreatic cancer is becoming increasingly deadly, with treatment options limited due to, among others, the complex tumor microenvironment (TME). This short communications study investigates pulsed low-dose-rate radiation (PLDR) as a potential alternative to conventional radiotherapy for pancreatic cancer neoadjuvant treatment.

Our ex vivo research demonstrates that PLDR, in combination with chemotherapy, promotes a shift from tumor-promoting to tumor-suppressing properties in a key component of the pancreatic cancer microenvironment we called CAFu (cancer-associated fibroblasts and self-generated extracellular matrix functional units). This beneficial effect translates to reduced desmoplasia (fibrous tumor expansion) and suggests PLDR’s potential to improve total neoadjuvant therapy effectiveness.

To comprehensively assess this functional shift, we developed the HOST-Factor, a single score integrating multiple biomarkers. This tool provides a more accurate picture of CAFu function compared to individual biomarkers and could be valuable for guiding and monitoring future therapeutic strategies.

Our findings support the ongoing NCT04452357 clinical trial testing PLDR safety and TME normalization potential in pancreatic cancer patients. The HOST-Factor will be used in samples collected from this trial to validate its potential as a key tool for personalized medicine in this aggressive disease.

---

Pancreatic cancer will soon become the second deadliest cancer in the USA^1^. Its most common type, pancreatic ductal adenocarcinoma (PDAC), is characterized by a significant stromal expansion, known as desmoplasia, which establishes a uniquely fibrous tumor microenvironment (TME).

PDAC-bearing pancreata encompass a distinctive TME, sustained by cancer-associated fibroblasts (CAFs) and self-generated extracellular matrix (ECM). These CAF/ECM functional units (CAF*u*) support PDAC cells, facilitate immunosuppression, and expand desmoplasia, via a forward vicious cycle driven by tumor-permissive (TP) CAF*u*-generated ECM^2-4^. Importantly, CAF*u* adopt both tumor-suppressive (TS) and TP functions^3^. The possibility of blocking CAF*u*’s TP functions while retaining the unit’s natural TS functions, triggering TME normalization, holds promise for novel therapeutic interventions^2, 3^.

Treatment of PDAC is challenging, with surgical resection being the only potential curative intervention. Total neoadjuvant therapy (TNT), combining chemotherapy and radiation prior to surgery, improves surgical outcomes^5^. Current standard PDAC chemoradiation regimens use radiation doses that serve primarily as an adjuvant treatment to surgery but are not sufficient on their own to eradicate this cancer. These doses are limited to prevent toxicities in the small bowel and stomach. Further, conventional chemoradiation can induce TP alterations in the TME where CAF*u* responses to therapy play a central role^6^. Hence, developing safer methods to escalate radiation dosages, without triggering TP-CAF*u* responses that further expand desmoplasia, could address the clinically unmet need for more effective TNT strategies. These methods, if proven safe, could also benefit future patients with locally advanced or metastatic PDAC.

Pulsed low-dose-rate radiation (PLDR), also known as pulsed-reduced-rate radiation, exhibits decreased toxicity compared to conventional radiotherapy (RT)^7^ and prevents RT-induced inflammation (*e*.*g*., lowers systemic levels of transforming growth factor beta –TGFβ)^8^. Since TGFβ-dependent factors and others promote TP-CAF*u* activation, and pro-tumoral desmoplastic expansion^4^, we investigated the effects of PLDR on CAF*u* traits and functions. For this, we used our well-known *ex vivo* human pancreatic CAF*u* culturing system (Supplementary Table – M&M *a & b*) to determine whether using PLDR, instead of conventional RT, could prevent the unwanted TP effects of TNT upon CAF*u* traits (Sup Figure 1A-F), and/or its desmoplastic expansion potential (SupFigure 1G-H).

**Figure 1.**
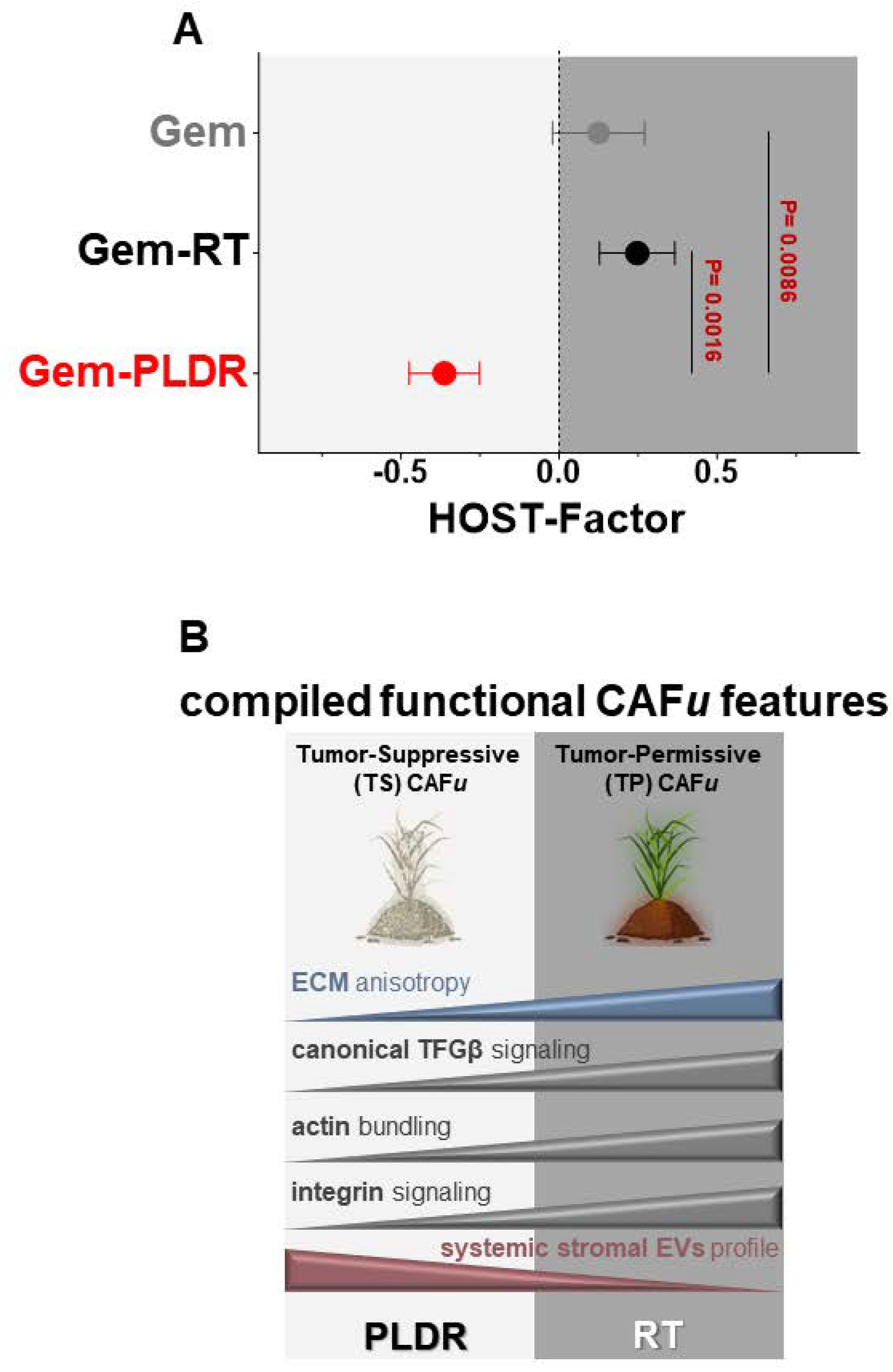
HOST-Factor: Quantifying CAF*u* function in response to TNT. **A-** Graph depicts the HOST-Factor values calculated from five functional assays employed to evaluate the impact of various neoadjuvant treatments on CAF*u* cultured using the 3D *ex vivo* system (see methods and references in the Supplementary Materials). Higher HOST-Factor values relative to gemcitabine (Gem) alone indicate a shift towards a more tumor-permissive (TP) phenotype in treated CAF*u*, while negative values suggest a tendency towards functional normalization (potential restoration of tumor-suppressive (TS) function, further supported by the functional assay in Supplementary 1H). Bullets represent mean ± standard error of the combined z-scores using the parameters measured in Supplementary 1B-F. **B-** Schematic diagram illustrating the individual tendencies of the measured values that constitute this study’s complied HOST-Factor. Selected elements from BioRender.com were incorporated. Note that the antibody-specific information was withheld as it relates to biomarkers that comprise the proprietary “HOST-Factor” signature.

Accurate assessment of CAF*u* functional states is essential for developing effective TME-targeted interventions. To address this, we commonly use our patient-derived *ex vivo* CAF*u* functional culturing system, which has enabled the identification of numerous biomarkers suggestive of CAF*u*’s TP/TS states (SupTable 1 – M&M a-c). However, relying on individual biomarkers can be imprecise due to an incomplete portrayal of CAF*u* function. For instance, αvβ5-integrin deficiency induces a TS CAF*u* with low TGFβ signaling, limited 3D-matrix adhesion activity, and an isotropic ECM that limits *de novo* activation of treatment-naïve CAFs^4^. Conversely, NetrinG1 knockdown in TP-CAF*u* triggers a complete functional TP-to-TS transition independent of altering ECM architecture^2^. Recognizing these complexities, we propose the <u>H</u>armonized <u>O</u>utput of <u>S</u>tromal <u>T</u>raits factor (HOST-Factor) – a single numerical score integrating a proprietary biomarker set data from several assays to comprehensively measure CAF*u*’s functional status (Figure 1).

Five established assays investigated the effects of gemcitabine (Gem), Gem-RT, and Gem-PLDR on treatment-naïve CAF*u* (SupFigure 1A-F). The assays measured:

1. ECM anisotropy: reflecting spatial organization of matrix fibers (SupTable – M&M d).
2. Canonical TGFβ signaling activity: a key driver of pro-tumorigenic fibrosis and immune tolerability (SupTable – M&M e).
3. Actin bundling modulation: influencing cellular contractility, ECM remodeling, and others (SupTable – M&M e).
4. Sustained activation of 3D-adhesions: indicative of integrin-dependent TP-CAF*u* function (SupTable – M&M e).
5. The presence of unique CAF*u*-generated extracellular vesicles: cell-cell communication mediators that can systemically inform on CAF*u*’s functional status (SupTable – M&M e & f).

*Ex-vivo* Gem-RT tended to nearly double the HOST-Factor score compared to Gem alone (P=0.5965). In contrast, Gem-PLDR had an opposite effect; 3-fold reduction compared to Gem-RT (P=0.0016), indicative of a TP-to-TS transition. This functional shift was confirmed in our desmoplastic expansion test (SupFigure 1G-H): Gem-RT treated CAF*u* ECM potently activated treatment-naïve fibroblasts (αSMA/F-actin=69.5%), while Gem-PLDR treated ECM had minimal impact (αSMA/F-actin=32.8%), demonstrating a significant (p<0.0001) 2-fold reduction in fibroblastic activation.

These data support the notion that Gem-PLDR supports a TP-to-TS transition in human pancreatic CAF*u*. Also, these findings validate the HOST-Factor as a comprehensive tool for quantifying CAF*u*’s function. Expanding the HOST-Factor to incorporate additional markers, reflecting the diverse cell types within the TME, could generate, in the future, even more informative scores, enhancing further the potential use of the HOST-Factor score for therapeutic guidance and monitoring purposes.

Our findings, aligned with reported studies^7-10^, lend strong support for NCT04452357, a PDAC phase 1 trial testing PLDR’s safety with escalated radiation doses and its TME normalization potential. Notably, the HOST-Factor will serve as a key TME functional assessment tool in this trial, with the goal of further validating its utility in guiding and monitoring therapeutic strategies.

## Abbreviations

(PDAC): Pancreatic ductal adenocarcinoma
(TME): tumor microenvironment
(CAF): cancer-associated fibroblast
(ECM): extracellular matrix
(CAF*u*): functional units composed of carcinoma-associated fibroblast and self-generated extracellular matrix
(TS): tumor-suppressive
(TP): tumor-permissive
(TNT): total neoadjuvant therapy
(PLDR): pulsed low-dose-rate radiation
(RT): radiotherapy
(TGFβ): transforming growth factor beta
(HOST-Factor): harmonized output of stromal traits factor
(Gem): gemcitabine

## Author contributions

Conceptualization and writing (EC, JEM, and JFB); data analyses and/or significant experimental contributions (TL, JFB, EH, KSR, JKW, DBVC, RF, JCG), discussions (JFB, TL, JEM, EH, and EC).

## Acknowledgments

We dedicate this work to Patricia Keely (pioneer in the field of ECM biology) and Neelima Shah who continues inspiring our research. This work was supported by Fox Chase institutional pilot funds intended to promote multidisciplinary translational efforts, the 5th AHEPA Cancer Research Foundation (EC), the Pancreatic Cancer Cure Foundation (EC), the Emerald foundation Inc. Black in Cancer Postdoctoral grant (JCG), NIH NCI grants: R01CA269660 (EC, JFB, and TL), 3R01CA269660-02S1 (EC), and U54CA272686 (EC), the equipment grant S10ODO23666, the DOD grant HT9425-23-1-0584 (EC and DBVC), and NIH/NCI P30CA06927 in support to the following facilities: Bio Sample Repository, Radiation, Biostatistics and Bioinformatics, Histopathology, Cell Culture, Imaging, and the Translational Protocols Laboratory.

## Supplemental Materials

**Supplementary Figure 1.**
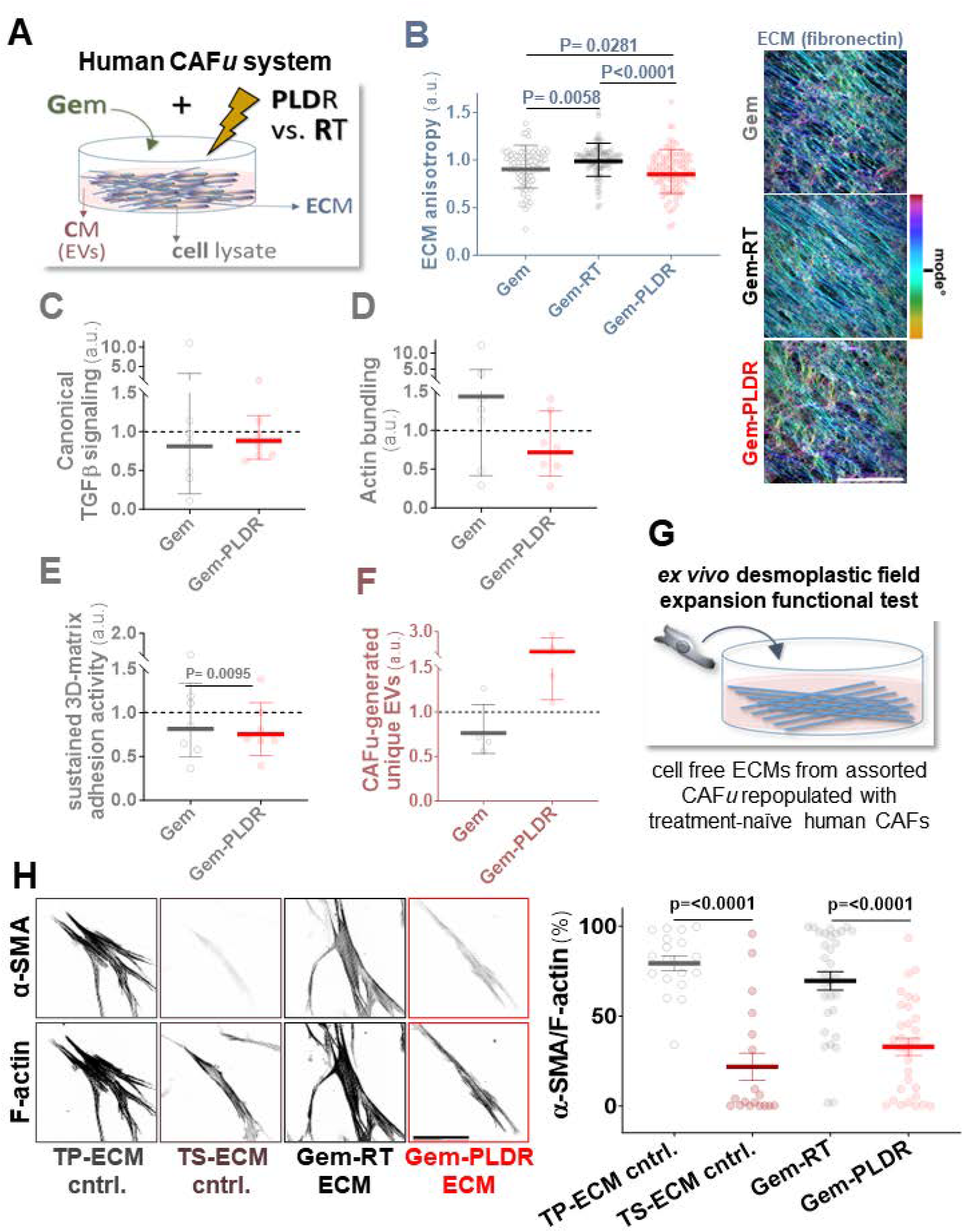
PLDR normalizes Human CAF*u* function. **A-** Schematic representation of *ex vivo* CAF*u* treatment conditions; gemcitabine alone (Gem), Gem plus conventional/continuous radiotherapy (RT), and Gem plus pulsed low-dose-rate radiation (PLDR). **B-** ECM anisotropy: Graph portraying the level of matrix fibers alignment (anisotropy) for each treatment group compared (normalized) to Gem-RT (set as 1.0 a.u.). Bullets represent anisotropy levels measured per image, averaged across 3 replicates and 4 human CAF lines. Larger numbers indicate increasing values of parallel fiber orientation. Pseudo-colored confocal images on the right show representative fiber angles, where cyan tone marks the mode angle and fibers deviated from this angle are represented by other colors. Scale bar = 100 µm. **C-E** Treatment-induced signaling changes: Graphs depict relative levels of key CAF*u* functional biomarkers compared to Gem-RT values (dotted line), gauged via immunoblot and normalized to GAPDH (loading control). **C-** Canonical TGFβ signaling reflecting both CAF*u*’s pro-fibrotic activity and potential for pro-tumoral immune tolerance. **D-** Actin bundling intensity (*e*.*g*., bundling protein detection), informing on the tumor permissive (TP) CAF*u*’s contractile ability, beyond α-SMA. **E-** 3D-matrix adhesion activity, informing on the reciprocal integrin-dependent cell-ECM signaling interactions that when sustained tend to render TP-CAF*u*. **F-** Presence of unique extracellular vesicles (EVs); elevated levels denote a shift towards a TS CAF*u* function. EVs were collected as published (SupTable – M&M e & f) and cargo was standardized using hexokinase as distinctive-EV loading control. Data in **C-F** represent geometric means ± geometric standard deviation across samples treated as indicated. **G-** *Ex vivo* desmoplastic field expansion functional verification: experimental schematic representation showing cell-free ECMs obtained from treated CAF*u* and repopulated with treatment-naïve human CAFs, overnight. **H-** Representative images and measured values showing percentage of α-SMA enriched stress fibers (F-actin), whereby ECMs produced by previously characterized TP-CAF*u* and from β5-integrin deficient TS-CAF*u*, respectively served as TP- and TS-ECM controls. Bar = 100 µm. Data points denote 1-3 cells per image and are represented as percentage means ± standard error of mean. This experiment was conducted in triplicate with 2 technical replicates per condition. Note that Gem-PLDR effectively phenocopied the function of control TS-ECMs preventing treatment-naïve CAFs from adopting an activated conformation, and thus limiting desmoplastic field expansion.

Please refer to the SupTable tabs “M&M” for information on published assay details, and “stats” for statistical details. Note: Antibody-specific information is withheld as it relates to biomarkers that comprise the proprietary “HOST-Factor”.

## Materials & Methods

### Ethical Statement

This study adhered to ethical guidelines, including those established by the Declaration of Helsinki, CIOMS, Belmont Report, and U.S. Common Rule. It was approved by the internal institutional review board at Fox Chase Cancer Center. Human specimens were collected with written informed consent for research, and personal information was safeguarded. Decoded samples, unlinked to the patients’ personal information, were provided to the researchers who harvested the fibroblastic cells used. All Fox Chase facilities used followed the International Society for Biological and Environmental Repositories and National Cancer Institute Best Practices for Bio-specimen Resources guidelines.

### CAF cell lines

Four heterogeneous human PDAC-derived CAF cell lines, obtained from untreated PDAC patients at time of surgery, were harvested, characterized, and cultured as functional CAF*u ex vivo*, according to our published methods (SupTable – M&M a & b).

### Gemcitabine and Radiation Treatments

Prior to total neoadjuvant treatment modalities, the effective IC50 of gemcitabine (Gem) (Pfizer; FCCC’s Pharmacy) effective for elimination of PDAC cells without compromising viability of CAF cells was established. For this, CAF and well-established human PDAC cells (Panc1) were cultured separately in the presence of increased Gem concentrations for five days. We found that 5.0 nM Gem effectively eliminated ∽50% of PDAC cells, with no significant reduction of CAF population (data not shown). CAF*u* were subjected to a daily treatment with Gem, either as a single agent, or in combination with gamma radiation during ECM production (Supplementary Table – M&M a& b). CAF*u* were treated with Gem alone or with Gem plus radiotherapy (RT); the results presented were for the conditions adjusted for *in vitro* settings using a total of 8 Gy per day. Radiation was administered using continuous RT or pulsed low-dose rate radiation (PLDR; the dose was fractionated into 40 pulses of 20 cGy bursts each, intercalated with periods of 3 min recovery, until 8 Gy were accumulated).

### Analysis of Therapeutic Response of CAF Units

All assays were conducted following our previously published protocols (for key references see SupTable tab “M&M”). Methods included CAF isolation, CAF*u* formation, characterization of CAF*u* using indirect immunofluorescence, immunoblotting, as well as EV isolation and characterization. Functional verification was done by repopulating the experimental acellular ECMs with treatment-naïve CAFs. Statistical data is in the “stats” tab of the Supplemental Table.

### HOST-Factor

The comprehensive response of CAF*u* to the assorted experimental conditions was obtained by standardizing the values of the selected CAF*u* traits presented in SupFigure 1B-F. A z-score was obtained by subtracting the mean value (µ), per trait set, from each corresponding data point (x), and divided by the standard deviation (σ) of the trait, as follows:

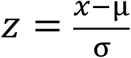

Z-scores from CAF*u*-generated EV levels were transformed to ensure for all HOST-Factor components equal biological significance and distinguish TP-CAF from TS-CAF characteristics. The HOST-Factor output indicated differences related to normalized values for each treatment. Positive outputs (>0) indicate enhanced tumor-promoting (TP) CAF*u* function, while negative values (<0) suggested that CAF*u* reduced TP qualities, gaining tumor-suppressive (TS) traits in response to the assorted treatments.

### Statistical analysis

Data were organized and calculated using Excel 2016 (Microsoft Office; SupTable - stats). Statistical significance was evaluated using Krustall-Wallis (one-way ANOVA) test, using GraphPad Prism 10 (V10.1.1) for Windows. Single CAF*u* parameters were graphed independently as geometric means with geometric standard deviations. Desmoplastic field expansion capabilities, and HOST-Factor scores are shown as means with standard errors of mean. Statistical significance between different conditions is indicated in the graphs when P value ≤ 0.05.

**Supplementary Table**

Tab M&M = list of studies reporting used methods.

Tab stats = calculated statistical values.

---

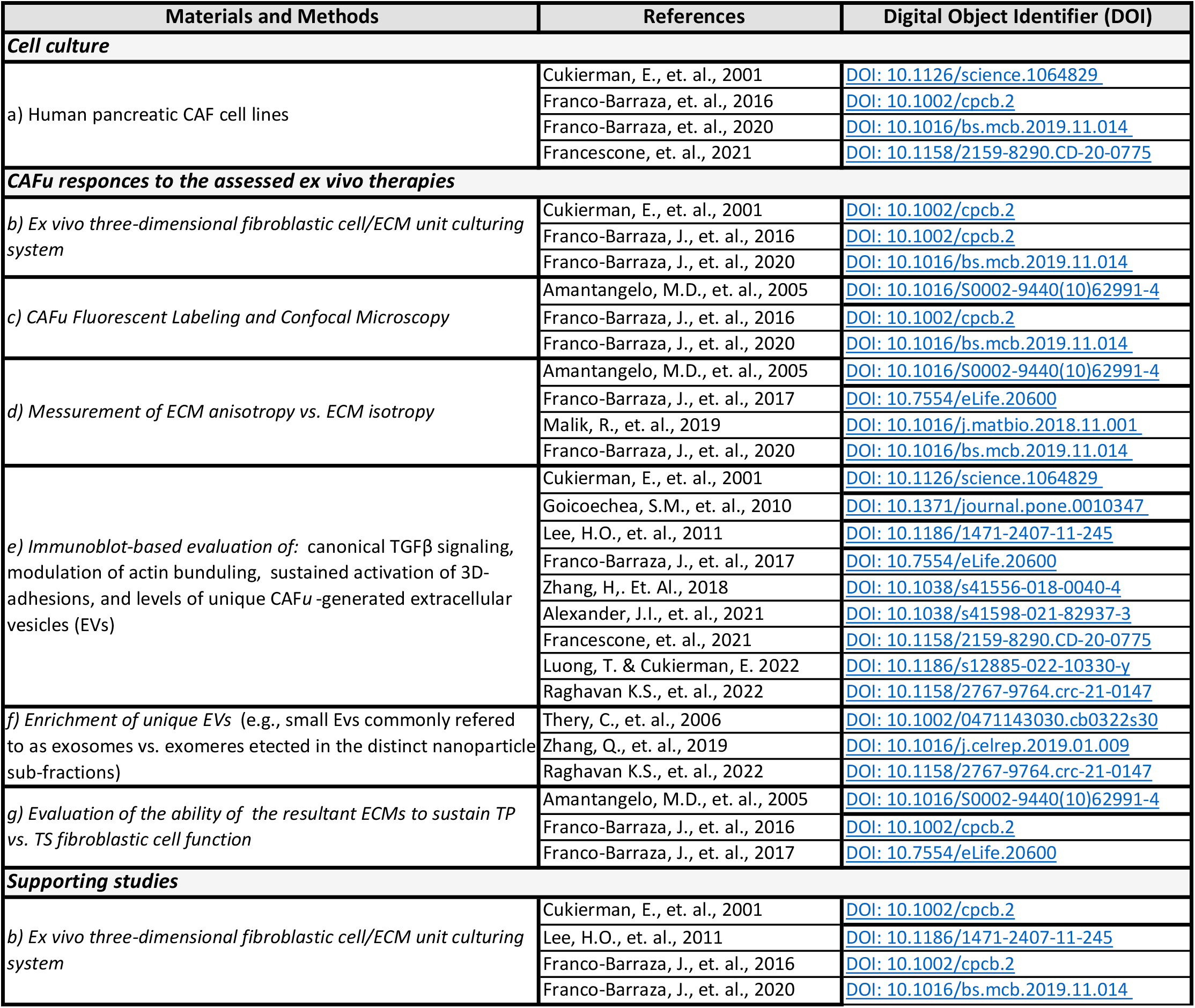
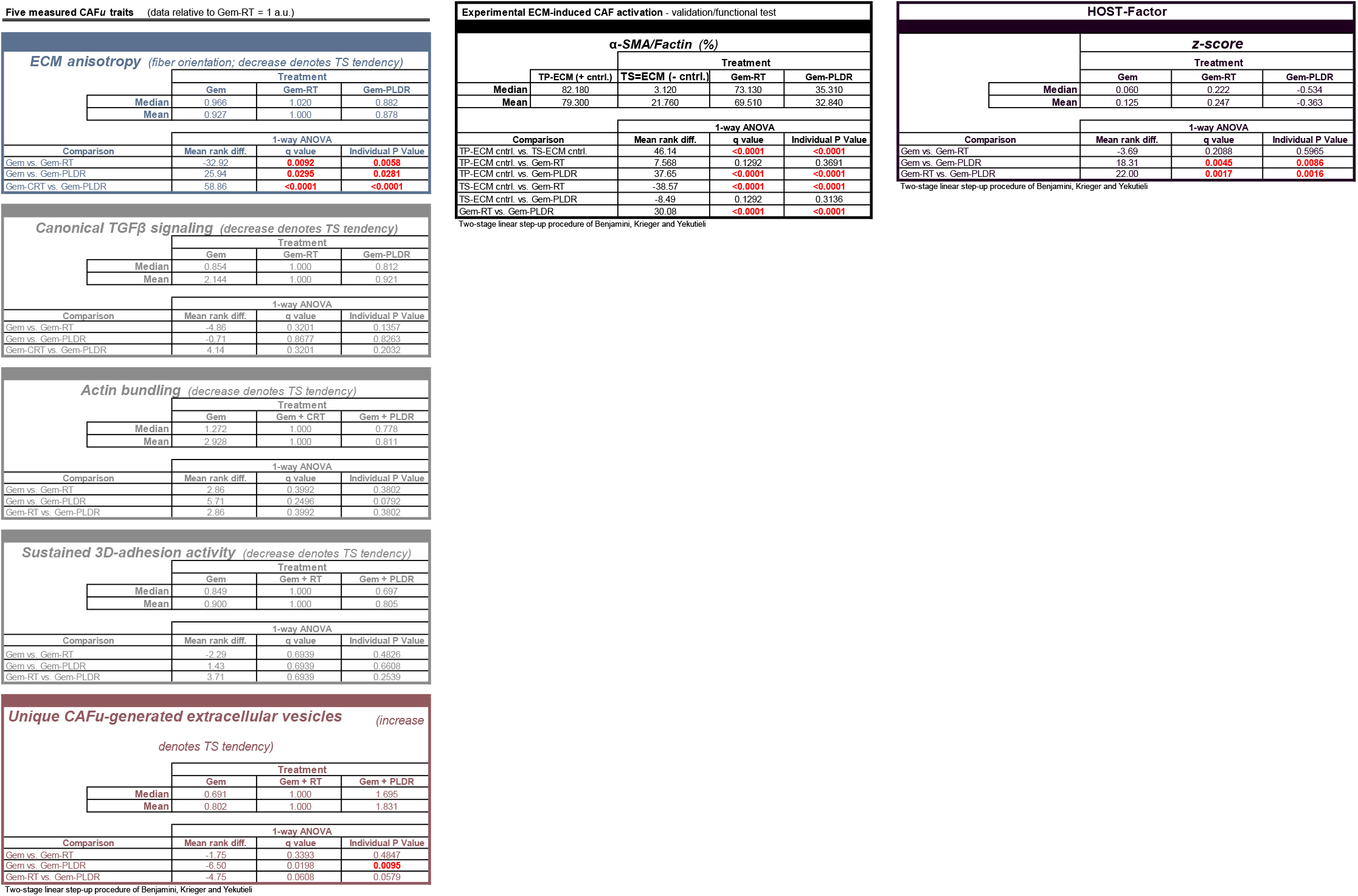

